# Analytical and Clinical Validity Study of FirstStep^Dx^ PLUS: A Chromosomal Microarray Optimized for Patients with Neurodevelopmental Conditions

**DOI:** 10.1101/083741

**Authors:** Charles H. Hensel, Rena Vanzo, Megan Martin, Sean Dixon, Christophe G. Lambert, Brynn Levy, Lesa Nelson, Andy Peiffer, Karen Ho, Moises Serrano, Sarah South, Kenneth Ward, E. Robert Wassman

## Abstract

**Introduction:** Chromosomal microarray analysis (CMA) is recognized as the first-tier test in the genetic evaluation of children with developmental delays, intellectual disabilities, congenital anomalies and autism spectrum disorders of unknown etiology.

**Array Design:** To optimize detection of clinically relevant copy number variants associated with these conditions, we designed a whole-genome microarray, FirstStep^Dx^ PLUS (FSDX). A set of 88,435 custom probes was added to the Affymetrix CytoScanHD platform targeting genomic regions strongly associated with these conditions. This combination of 2,784,985 total probes results in the highest probe coverage and clinical yield for these disorders.

**Results and Discussion:** Clinical testing of this patient population is validated on DNA from either non-invasive buccal swabs or traditional blood samples. In this report we provide data demonstrating the analytic and clinical validity of FSDX and provide an overview of results from the first 7,570 consecutive patients tested clinically. We further demonstrate that buccal sampling is an effective method of obtaining DNA samples, which may provide improved results compared to traditional blood sampling for patients with neurodevelopmental disorders who exhibit somatic mosaicism.

**Clinical scenario:** Neurodevelopmental disabilities, including developmental delays (DD), intellectual disabilities (ID), and autism spectrum disorders (ASD), affect up to 15% of children (1). In the majority of cases, a child’s clinical presentation does not allow for a definitive etiological diagnosis. In such cases, CMA is recommended as the first-tier test that should be used to evaluate for a potential genetic etiology (2–7). A definitive genetic diagnosis allows patients to more often receive appropriate medical care tailored to their condition, as reflected by medical management changes and improved access to necessary support and educational services (8–13).

**Test description:** FirstStep^Dx^ PLUS (FSDX) is an optimized clinical microarray test provided in the context of a comprehensive clinical service. Testing starts with either a non-invasive buccal swab sample or traditional blood sample from which DNA extraction using a Gentra Puregene^®^ kit specific to the sample type (Qiagen, Inc., Valencia, CA) is performed in one of several contracted CLIA/CAP credentialed laboratories according to manufacturers’ protocols. High quality genomic DNA is fragmented, labeled and hybridized to FSDX arrays using reagents, equipment and methodology as specified by by the manufacturer (Affymetrix, Inc., Santa Clara, CA)(14). Washed arrays are scanned and raw data files are processed to CYCHP files using a reference file comprising at least 100 samples with normal array findings. Data analysis is performed using Chromosome Analysis Suite software version 2.0.1 (Affymetrix). Hybridization of patient DNA to oligonucleotide and SNP probes is independently compared against a previously analyzed cohort of normal samples to call CNVs and allele genotypes. The percentage mosaicism of whole-chromosome aneuploidies is determined using the average log_2_ ratio of the entire chromosome (14).

**Microarray design:** FSDX was optimized by the addition of 88,435 custom probes targeting genomic regions strongly associated with ID/DD/ASD (15–24). This was effected, under GMP by Affymetrix, to the CytoScanHD platform using their microarray design process specifications which have been previously described (14). This is consistent with the ACMG recommendation of “enrichment of probes targeting dosage-sensitive genes known to result in phenotypes consistent with common indications for a genomic screen” (25). Critical regions that did not meet a desired probe density ≥1 probe/1000 bp on the CytoScanHD were supplemented with additional probe content to allow for improved detection of smaller deletions and duplications in these critical regions. Finally, additional probes were added to improve detection of CNVs encompassing genes involved in other well-characterized neurodevelopmental disorders, for example *GAMT* (26) and *GATM* (27). All incremental probes were added in substitution for probes deemed sub-optimal by Affymetrix and previously masked, bringing FSDX to a grand total of 2,784,985 probes. Custom SNP probes (n =416) on FSDX are targeted by 12 oligonucleotides, three for each strand of each allele, which is approximately double the typical probe coverage for SNPs.

**Test interpretation:** CYCHP files are evaluated by ABMG certified cytogeneticists. Determination of CNVs is consistent with established cytogenetic standards. A minimum of 25-consecutive impacted probes is the baseline determinant for deletions and 50 probes for duplications independent of variant size. Rare CNVs are determined to be "pathogenic" if there is sufficient evidence published (at least two independent publications) to indicate that haploinsufficiency or triplosensitivity of the region or gene(s) involved is causative of clinical features or of sufficient overall size (28). If however, there is insufficient but at least preliminary evidence for a causative role for the region or gene(s) therein they are classified as variants of unknown significance (VOUS) independent of CNV size. Areas of absence of heterozygosity (AOH) are also classed as VOUS if of sufficient size and location to increase the risk for conditions with autosomal recessive inheritance or conditions with parent-of-origin/imprinting effects. Other CNVs are typically not reported after determination that they most likely represent normal common population variants and are contained in databases documenting presumptively benign CNVs, e.g., the Database of Genomic Variants (DGV) (29). These parameters were standard independent of the microarray used for analysis in comparative studies.

**Public health importance:** A definitive genetic diagnosis facilitates patient access to appropriate and necessary medical and support services. Defining the underlying genetic cause of DD/ID/ASD and/or multiple congenital anomalies (MCA) in each unique patient is vital to understanding etiology, prognosis, and course. It informs physicians of potential comorbid conditions for which a patient should be evaluated and treated proactively and optimally. Improved understandings of the appropriate therapeutic and behavioral approaches to that patient are also enabled. Genetic testing is best provided in the context of an integrated service (30), so FSDX provides comprehensive, clear, readable, and personalized reports for the healthcare provider and a family-friendly section to facilitate understanding of the often-complex results. The report is complemented by availability of pre- and post-test genetic counseling and technical support to providers. Moreover, FSDX includes personalized insurance pre-authorization and appeals assistance to overcome barriers encountered by both providers and families that, in many circumstances, prevent access to crucial genetic testing services (10–11).

**Published reviews, recommendations and guidelines:** The American College of Medical Genetics (ACMG) (2,3), the American Academy of Child and Adolescent Psychiatry (4,7), the American Academy of Pediatrics (5), and the American Academy of Neurology/Child Neurology Society (6) recommend CMA as the first-tier test in the genetic evaluation of children with unexplained DD, ID, or ASD. Considerable data supporting these guidelines are documented in numerous reviews and publications (31–36). The ACMG has also published guidelines on both array design (25) and the validation of arrays, including validation of a new version of a platform in use by the laboratory from the same manufacturer, and of additional sample types (28).

## EVIDENCE OVERVIEW

### Validation of novel (Blood vs. Buccal) sample type on the established and optimized platforms

It is highly desirable to avail clinicians and families a less invasive sampling method for individuals with ID/DD/ASD due to the potential implications of venipuncture in many such individuals. We validated a buccal sampling methodology in conjunction with two independent CLIA-certified laboratories, ARUP (Salt Lake City, UT) and Fullerton Genetics Center (Asheville, NC) first on the CytoScanHD array and then the FSDX array. After preliminary consideration of multiple saliva and buccal DNA collection kits, we selected the ORAcollect-100^®^ (DNA Genotek, Ottawa)(now cleared as a class II IVD medical device: ORAcollect-DX^®^) based upon ease of use for the intended patient population, ease of shipping processes, DNA stability, and post-extraction quality studies for further validation. Buccal swabs and blood samples from twenty-two individuals underwent parallel microarray analysis for concordance by each protocol. Twenty-two individuals’ buccal and blood samples were analyzed in terms of array quality control metrics and no significant difference between the two sample sources was observed. In addition, there was 100% concordance of CNV calls between the two sample types. Finally, twenty-three individuals’ blood and buccal samples underwent microarray analysis on the FSDX platform. Again no significant difference between the two sample sources in terms of quality metrics and 100% clinical concordance of CNV calls was observed.

### Clinical Validation of a new version of a previously established platform

FSDX was validated independently in four CLIA-certified laboratories (Asuragen – Asuragen, Inc., Austin, TX; AGI – Affiliated Genetics, Inc., Salt Lake City, UT; Fullerton - Mission Hospital/Fullerton Genetics Center, Asheville, NC; CUMC – Columbia University Medical Center, Dept. of Pathology and Cell Biology, New York, NY.) all previously familiar with performing CMA on CytoScanHD (or its predecessor the Affymetrix 2.7M Cytogenetics array) and cross-validated between these laboratories as well. Data demonstrating the concordance of FSDX with these alternative arrays and across laboratories are shown in Tables I–II. A total of six samples from patients having clinically significant CNVs findings on prior clinical analysis were re-analyzed using FSDX. This study mirrors ACMG guidance on clinical validation of a new version of a previously established platform when the total probe content change is less than 5% (here 3.3% increase) (28). There was complete clinical concordance between the initial clinical results and the results generated with FSDX in two independent laboratories (AGI and CUMC). Although the cross-platform and cross-laboratory results are unequivocally clinically concordant, minor differences in breakpoint determinations were observed in the majority of analyses (8 of 11), which is expectable and reasonable within the limits of the technology and interpretation overall. The differences in breakpoints between FSDX and CytoScanHD resulted in CNVs that were smaller, but only on average by ≤0.25% of the total CNV size. It was previously observed that arrays with increasing increased probe density result in smaller estimates of total CNV size presumably through the higher resolving power with increased probe density (14). In contrast, only a single inter-laboratory analysis of FSDX differed in breakpoint calls by the different cytogeneticists, and in this case the change was only 0.08% of the overall CNV size.

**Table I:** Clinical validity of the FirstStep^Dx^ PLUS array. Samples from patients analyzed clinically on commercially available Affymetrix microarrays were run independently on FSDX. Laboratory designations are as follows: Asuragen – Asuragen, Inc., Austin, TX; AGI – Affiliated Genetics, Inc., Salt Lake City, UT; Fullerton - Mission Hospital/Fullerton Genetics Center, Asheville, NC; CUMC – Columbia University Medical Center, Dept. of Pathology and Cell Biology, New York, NY. All arrays used clinically were purchased from Affymetrix, Inc., Santa Clara, CA.

| Gender | Clinical Laboratory | Clinical Array | Clinical Result | Validation Laboratory 1 | Validation Result 1 | Custom Probes | Validation Laboratory 2 | Validation Result 2 |
| --- | --- | --- | --- | --- | --- | --- | --- | --- |
| M | ARUP | 2.7M Array | arr 15q11.2q14(18,446,422-32,204,913)x2hms; 15q25.1qter(77,139,194-100,201,798)x2hms (hg18) | AGI | arr 15q11.2q14(22,752,398-34,652,683)hms; 15q25.1q26.3(79,465,893-102,429,049)hms(hg19) | 1191 | CUMC | arr 15 q11.2q14(22,752,398-34,652,683)x2hms; 15q15.1q21.1(42,484,889-46,012,198)x2hms; 15q25.1 q26.3(79,465,893-102,429,049)x2hms |
| M | ARUP | CytoScanHD | arr 16p11.2(29,580,020-30,178,404)x3 (hg19) | AGI | arr 16p11.2(29,421,694-30,177,916)x3(hg19) | 0 | CUMC | arr 16p11.2(29,580,020-30,178,406 x3)(hg19) |
| M | Asuragen | CytoScanHD | arr (1-22)x2,(XY)x1 (hg19) - LCSI on CHR1 | AGI | arr (1-22)x2,(XY)x1 (hg19) - LCSI on CHR1 (10.5 Mb 1p) |  | CUMC | arr (1-22)x2,(XY)x1 (hg19)- LCSI on CHR1 |
| F | Fullerton | CytoScanHD | arr 16p12.2(22,086,845-22,442,007)x1 (hg19) | AGI | arr 16p12.2(21,946,522-22,442,007)x1 (hg19) | 0 | CUMC | arr 16p12.2(22,086,845-22,442,007)x1 (hg19) |
| F | Fullerton | CytoScanHD | arr 22q13.33(49,423,616-51,197,838)x1 (hg19) | AGI | arr 22q13.33(49,424,409-51,197,838)x1(hg19) | 56 | CUMC | arr 22q13.33(49,423,616-51,197,838)x1(hg19) |
| M | Asuragen | CytoScanHD | arr 15q11.2q13.1(23,620,191-28,522,838)x1 (hg19) | AGI | arr 15q11.2q13.1(23,642,700-28,561,671)x1(hg19) | 0 |  |  |

**Table II:**
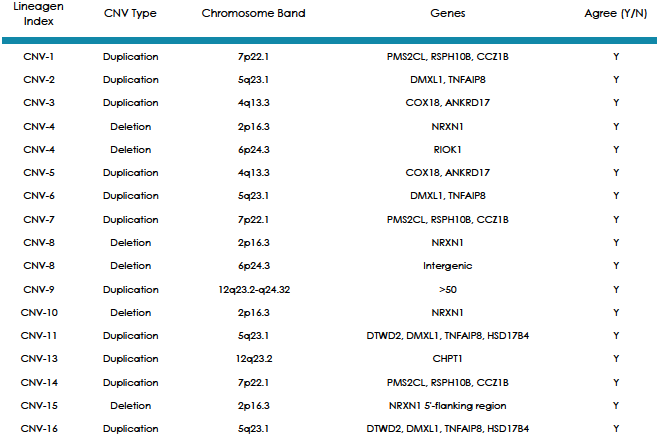
Research findings validated on FirstStep^Dx^ PLUS. Samples derived from research studies with significant CNVs present (15) which spanned both custom and standard probes were evaluated independently on FSDX in two laboratories and evaluated for agreement with the research findings as well their inter-laboratory concordance.

| Lineagen Index | CNV Type | Chromosome Band | Genes | Agree (Y/N) |
| --- | --- | --- | --- | --- |
| CNV-1 | Duplication | 7p22.1 | PMS2CL, RSPH10B, CCZ1B | Y |
| CNV-2 | Duplication | 5q23.1 | DMXL1, TNFAIP8 | Y |
| CNV-3 | Duplication | 4q13.3 | COX18, ANKRD17 | Y |
| CNV-4 | Deletion | 2p16.3 | NRXN1 | Y |
| CNV-4 | Deletion | 6p24.3 | RIOK1 | Y |
| CNV-5 | Duplication | 4q13.3 | COX18, ANKRD17 | Y |
| CNV-6 | Duplication | 5q23.1 | DMXL1, TNFAIP8 | Y |
| CNV-7 | Duplication | 7p22.1 | PMS2CL, RSPH10B, CCZ1B | Y |
| CNV-8 | Deletion | 2p16.3 | NRXN1 | Y |
| CNV-8 | Deletion | 6p24.3 | Intergenic | Y |
| CNV-9 | Duplication | 12q23.2-q24.32 | >50 | Y |
| CNV-10 | Deletion | 2p16.3 | NRXN1 | Y |
| CNV-11 | Duplication | 5q23.1 | DTWD2, DMXL1, TNFAIP8, HSD17B4 | Y |
| CNV-13 | Duplication | 12q23.2 | CHPT1 | Y |
| CNV-14 | Duplication | 7p22.1 | PMS2CL, RSPH10B, CCZ1B | Y |
| CNV-15 | Deletion | 2p16.3 | NRXN1 5'-flanking region | Y |
| CNV-16 | Duplication | 5q23.1 | DTWD2, DMXL1, TNFAIP8, HSD17B4 | Y |

As further demonstration of the clinical validity of this array, 16 samples from an earlier research study using an alternative technology platform (Illumina, San Diego, CA) (15) were analyzed on FSDX in two participating CLIA-laboratories: ARUP &Asuragen (Table II). Samples all had significant CNVs present, which had been validated by quantitative PCR in the research study, and spanned both custom and standard probes. All results were concordant both across platforms and between laboratories.

### Inter-laboratory clinical performance validation

Further evidence of the inter-laboratory performance (Table III) is shown on twelve additional patients with clinically significant CNVs detected by clinical testing with FSDX at Asuragen, and then re-analyzed by both AGI and CUMC, again with completely concordant results. In addition, two patients run clinically at CUMC were concordant with results subsequently generated by AGI, and six patients run clinically at AGI were concordant with results generated by Fullerton Genetics Laboratory Center.

**Table III:**
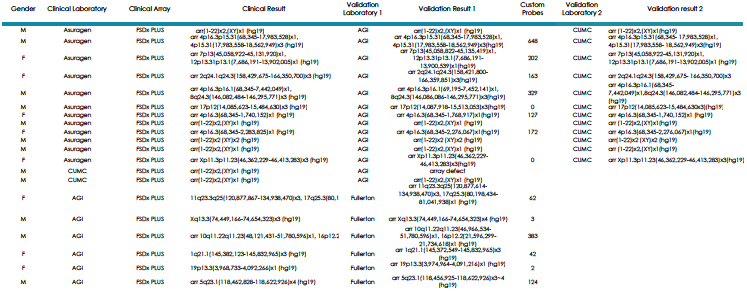
Inter-laboratory validation of the FirstStep^Dx^ PLUS array. Independent samples from patients analyzed clinically at our participating laboratories were re-analyzed at other participating laboratories. Laboratory designations are as described in Table II.

These data clearly demonstrate the ability of FSDX to detect copy number variants on a consistent basis in independent laboratories, across a range of genomic locations with all samples yielding concordant results to those performed on three separate array platforms. The excellent performance in cases with CNVs previously detected in regions with custom probes supports the overall clinical consistency and appropriate performance of the custom probe content on the array.

### Extended comparative clinical sensitivity studies in a in real-world clinical population

Data from 7570 consecutive patient samples tested clinically with FSDX from July 2012 (when the optimized FirstStep^Dx^ PLUS (FSDX) microarray was implemented into routine use) through May 2016 are shown in Figure 1. Overall there were 717 (9.5%) pathogenic abnormalities and 1534 (20.2%) VOUS observed, or a 29.7% overall CNV diagnostic yield for potentially abnormal findings. We also compared these results to patients who were tested by Lineagen through the same referral channels and in comparable patient cohorts and general time windows using the 2.7M and CytoScanHD arrays. The 2.7M arrays (n=378) detected 7.4% pathogenic and 8.5% VOUS for an overall yield of 15.9%. The higher density CytoScanHD arrays (n=1194) detected 9.0% pathogenic and 14.2% VOUS for an overall yield of 23.2%. It is clear that these incremental probe additions translate into potentially clinically significant yields, likely with resultant improvements in patient medical management.

**Figure 1:**
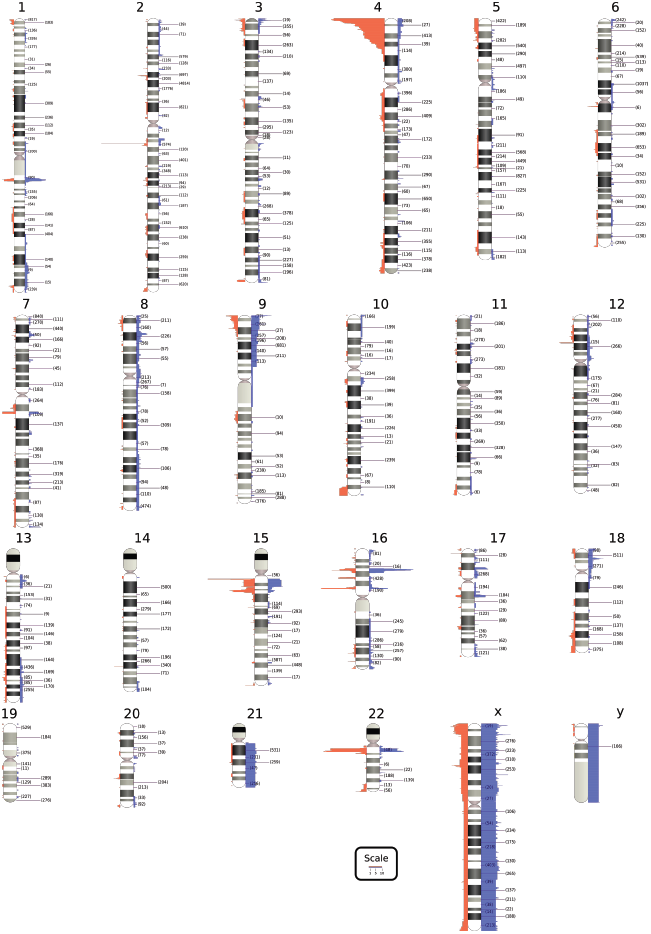
Summary of clinical CNV findings using FirstStep^Dx^ PLUS. Reported clinical findings are displayed next to chromosome ideograms. Deletions are shown on the left in red, and duplications are shown on the right in blue. Numbers in parentheses after chromosome band labels indicate the number of custom-designed probes in those bands. An excess of abnormalities is observed in the 4p region due to a research cohort; however this data is not reflected in the overall detection rates cited in the text.

Figure 1 visually summarizes the genomic range of CNVs detected to date using FSDX. These include known microdeletion and microduplication syndrome regions as well as variants of unknown significance.

### Quality Control

Similar to the CytoScanHD, the independent analysis of CNVs with oligonucleotide and SNP probes provides both detection and confirmation simultaneously (14). Three empirically determined quality control metrics are used that reflect overall data quality on an Affymetrix array: 1) waviness-SD, 2) median of the absolute values of all pairwise differences (MAPD), and 3) SNPQC (measure of how well genotype alleles are resolved). The criteria determined empirically for the CytoScanHD array extend to the FSDX array by virtue of design, with all three needed to meet minimum requirements for an array to be analyzed. FSDX arrays meeting these requirements can be interpreted over 99% of the time.

### Analytic validity

To rigorously demonstrate the comparable analytic validity of FSDX, we employed an independent tool, the Golden Helix, Inc. Copy Number Analysis Method (CNAM) (Golden Helix, Inc. Bozeman, MT)(36).First, Affymetrix Chromosome Analysis Suite (ChAS) version 2.1 (Affymetrix, Santa Clara, CA) was used to create an evaluation set of 205 samples with no reportable clinical finding from a single laboratory. To characterize the analytical function of custom probes added to the FSDX microarray, we used the sum of raw probe signal data across the array as a proxy for input DNA concentration. Across the 205 samples, there was roughly a 4.6 fold difference in total array signal. We then plotted individual probe signal intensity vs. total array signal for all probes on the array and used regression analysis to calculate slopes and y-intercepts for each probe. We divided total intensity by 933,696,453 (the lowest total intensity sum) to generate an X-axis that ranged from 1 to 4.6. Finally, we plotted y-intercept vs. slope for each probe and compared custom probes to CytoScanHD probes. Figure 2 shows a scatter-plot based on data from this subset of 205 samples. Data in red reflect custom probes only found on FSDX, while data in black designate standard CytoScanHD probes. The plot indicates that there is significant overlap between the custom probes and standard probes. However, at the extreme, a cluster of custom probes, with slope and intercept near zero, do not appear to respond well to increasing DNA input. Thus, these are less likely to be strongly copy number responsive, but are not necessarily non-informative. This “sub-par” population represents approximately 20% of the custom FSDX probes. This shift in performance characteristics of the single-pass design custom probes is still within the overall desired operating parameters; it suggests at most a 0.64% (% poorly functioning probes over total probe content) deviation in overall analytical sensitivity. However, given that the total probe content on FSDX is 3.27% greater, these data suggest a net gain in sensitivity of 2.63% (net increase in total optimally performing probes compared to CytoScanHD).

**Figure 2:**
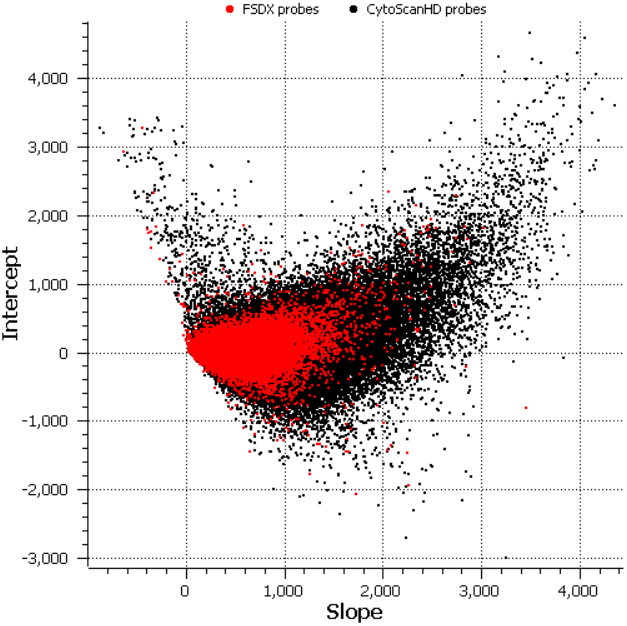
Functional Overlap of Custom Probes with CytoScanHD Probes. Histogram of slope and Y- intercept for each probe on the FirstStep^Dx^PLUS array. Probes shown in red are custom probes found only on FSDX, which overlie probes shown in black common to both FSDX and CytoScanHD. Probes with slope and intercept values near 0 are considered to be sub-optimal probes.

We next analyzed the relative impact of the custom probes compared to the standard array probes on overall detection rates of CNVs. The weighted log_2_ ratios from 184 arrays that each had at least one clinically reported finding previously analyzed using ChAS in a single laboratory were evaluated again with CNAM (36). This utilized the univariate option with no moving window; a maximum of 10 segments per 10,000 probes; a minimum segment size of 1 marker; and a stringent permutation p-value threshold of 0.001. After segmentation, we classified segments as losses if their mean was <0.20 with 25 or more probes and classified segments as gains if their mean was >0.20 with 50 or more probes (consistent with our clinical workflows). These calls were compared to the clinically reported findings generated with ChAS. The analysis with CNAM was performed in two ways. First, only probes present on the CytoScanHD standard array were considered. Second, FSDX custom probes in addition to the standard probes were considered. All clinically relevant findings were observed using both analysis methods (Table IV). Each reported CNV was detectable independent of the presence of custom probes in the CNV and independent of the use of the custom probes in the analysis. These data show no evidence that the previously described sub-optimally responding subset of custom probes interfere with the function of responsive probes, whether standard or custom, or with the overall sensitivity for CNV detection.

**Table IV:** Analytic validity of FirstStep^Dx^ PLUS. Data from clinical samples were evaluated using CNAM on weighted log_2_ ratios from 184 arrays as described above. The data from CNVs with and without custom FSDX probe content were evaluated to determine any discrepancy in detection based on inclusion of the custom probes and no evidence of non-concordance was observed.

| Chr | # Findings |  |  |  | # findings with Custom Probes | Agreement +/- custom probes |
| --- | --- | --- | --- | --- | --- | --- |
|  | Abnormal |  | VOUS |  |  |  |
|  | Deletions | Duplications | Deletions | Duplications |  |  |
| chr1 | 1 |  | 2 | 4 | 1 | Y |
| chr2 | 2 | 1 | 5 | 7 | 9 | Y |
| chr3 |  | 2 | 4 | 2 | 1 | Y |
| chr4 | 2 |  |  | 5 | 3 | Y |
| chr5 | 1 | 1 | 4 | 2 | 3 | Y |
| chr6 |  |  | 6 | 5 | 2 | Y |
| chr7 | 2 |  | 4 | 5 | 0 | Y |
| chr8 | 1 |  | 2 | 1 | 2 | Y |
| chr9 | 1 |  | 6 | 7 | 5 | Y |
| chr10 |  |  | 3 | 4 | 3 | Y |
| chr11 |  | 2 |  | 3 | 1 | Y |
| chr12 |  | 2 | 1 | 3 | 3 | Y |
| chr13 | 3 |  | 1 | 1 | 3 | Y |
| chr14 |  |  | 3 | 2 | 0 | Y |
| chr15 | 5 | 1 | 9 | 2 | 1 | Y |
| chr16 | 5 | 3 | 4 | 5 | 5 | Y |
| chr17 | 1 | 4 | 3 |  | 1 | Y |
| chr18 | 1 |  | 3 | 2 | 2 | Y |
| chr19 |  |  | 1 |  | 0 | Y |
| chr20 |  |  | 2 | 1 | 0 | Y |
| chr21 |  | 1 |  | 1 | 1 | Y |
| chr22 | 5 | 1 | 1 |  | 6 | Y |

The data in Table I also demonstrate the analytic validity of custom probes that are included in some of the clinically reported findings. Increased sensitivity and resolution has been demonstrated with increasing probe density (14), and since the custom probes were added specifically to regions important for ASD and other neurodevelopmental disorders, the resolution of FSDX for these conditions is predicted to be enhanced, since approximately 80% of the custom probes are fully analytically responsive to DNA input. Preliminary analysis (data not shown) suggests that the overall detection of CNVs may be increased by inclusion of the custom probes in this analysis, and further evaluation of this is under investigation.

### Clinical utility

The goal of any diagnostic test is improvement in medical management of patients. This is achieved first by reaching the correct diagnosis and second by following the appropriate management & surveillance procedures for that diagnosis. CMA has a clear and positive impact on medical management, as documented in several studies (8–13). Further, CMA testing often results in a correct,additional, or modified diagnosis for those who are difficult to diagnose with clinical observation alone (37, 38). Even in patients with well-defined conditions, better determining CNV breakpoints with a higher resolution methodology can provide information beyond what is known or assumed from other tests (39). A test, like CMA, which allows us to identify multiple individually rare diagnoses is difficult to assess by traditional measures of clinical utility. This is due, in part, to the fact that each potential diagnosis has specific benefits in terms health and cost of care for a given patient. However, the literature has documented numerous improvements in care (8–13) as a result of reaching a genetic diagnosis for individuals with DD, ID, and/or ASD. Further this represents this a continuum as clinical understanding and care evolves stepwise as a direct result of improved diagnostic clarity. The ability of increased probe density in genomic regions of interest to improve diagnosis is becoming apparent, and each additional correct diagnosis allows incremental opportunities for increased composite clinical utility of such tools.

### Limitations

Given the rarity of individual CNVs and current limits of understanding, this is a relatively small clinical validation cohort. This could limit the certainty and range of conclusions from this evaluation, however it exceeds established guidelines for such validations (28). Analytical validity across millions of data points as in a CMA, can only be assessed by in silico methods such as applied here, however this is superior to mere presumptions of performance typical in the literature.

CMA has proven an important clinical diagnostic advance, however it is limited by the failure to detect a diagnosis in still a majority of affected individuals even with the increased performance with added probe content in FSDX. However, emerging evidence suggests that another significant portion of these cases, which is negative on CMA will have mutations detectable by massively parallel sequencing (NGS) in the future complementing CMA (41). In addition, roughly two-thirds of reported CNVs are VOUS, largely due to the limitations of our clinical experience with these rare conditions. The use of new informatics approaches and databases are emerging to better define the potential relevance and pathogenesis of such currently uncertain findings (42).

## Conclusions

Analytic validation of FirstStep^Dx^ PLUS coupled with over three years of clinical use have demonstrated the utility of FirstStep^Dx^ PLUS in identifying genetic causes for neurodevelopmental conditions. The simple non-invasive sampling procedure, high clinical sensitivity and extensive support services make FirstStep^Dx^ PLUS an ideal choice for the clinical genetic evaluation of patients with these disorders.

## Ethics Statement

All involvement with human subjects was done in accordance with the World Medical Association <u>Declaration of Helsinki</u>, and with patient informed consent for anonymous post-clinical testing use of specimens and data.

## Acknowledgements

We thank Kenny Lentz for database management and support, Porter Rickabaugh for development of the FSDX diagnoses ideogram, and the entire Lineagen team for their remarkable energy and dedication to those with neurodevelopmental disabilities of the entire Lineagen team. We also thank the professional and laboratory staff of the laboratories that participated in work.

